# Non-invasive neuromodulation using rTMS and the Electromagnetic-Perceptive Gene (EPG) facilitates plasticity after nerve injury

**DOI:** 10.1101/851444

**Authors:** Carolina Cywiak, Ryan C. Ashbaugh, Abigael C. Metto, Lalita Udpa, Chunqi Qian, Assaf A. Gilad, Ming Zhong, Galit Pelled

**Affiliations:** Department of Biomedical Engineering, Michigan State University, East Lansing, MI, USA; The Institute of Quantitative Health Science and Engineering, Michigan State University, East Lansing, MI, USA; Department of Electrical and Computer Engineering, Michigan State University, East Lansing, MI, USA; Department of Radiology, Michigan State University, East Lansing, MI, USA

## Abstract

Peripheral nerve injury leads to altered cortical excitation-inhibition balance which is associated with sensory dysfunctions. We tested if non-invasive repetitive transcranial magnetic stimulation (rTMS) which has shown to induce neuronal excitability, and cell-specific magnetic activation via the Electromagnetic-perceptive gene (EPG) which is a novel gene that was identified and cloned from *Kryptopterrus bicirrhis* and demonstrated to evoke neural responses when magnetically stimulated, can restore cortical excitability. A battery of behavioral tests, fMRI and immunochemistry were performed in the weeks following limb denervation in rats. The results demonstrate that neuromodulation significantly improved long-term mobility, decreased anxiety and enhanced neuroplasticity. The study also identifies the acute post-injury phase as a critical time for intervention. Moreover, the results implicate EPG as an effective cell-specific neuromodulation approach. Together, these results reinforce the growing amount of evidence from human and animal studies that are establishing neuromodulation as an effective strategy to promote plasticity and rehabilitation.

## Introduction

Twenty million individuals in the United States are suffering from peripheral nerve injury. Current strategies to facilitate recovery following peripheral nerve injury mainly focus on manipulating the activity of the injured and the non-injured limbs. However, in spite of refined surgical techniques and the available rehabilitation strategies, the clinical outcome in adults is generally poor with persisting sensory dysfunction and pain complications^1^.

Over the last 35 years, studies have shown that acute or chronic disturbance to sensory afferents is reflected in distorted cortical representations. These anatomical and functional changes are evident by electrophysiology and fMRI methods and may impact clinical outcome. For example, human studies suggest a strong correlation between abnormal post-injury cortical responses that are often observed with fMRI to the degree of sensory dysfunctions and phantom limb pain^2-4^. The neural mechanisms implicated in the post-injury cortical changes have been extensively studied in animal models; Studies indicate that peripheral injury evokes cellular mechanisms effecting immediate^5^ and long-term^6-9^ function of the primary somatosensory cortex (S1) contralateral and ipsilateral to the injured limb. These mechanisms include alteration in the excitation-inhibition balance^10^, changes in GABAergic function^11^, and increases in the activity of inhibitory interneurons in cortical layer 5 (L5) in the affected (deprived) cortex^12-14^. Therefore, it is conceivable that post-injury cellular changes affect neurorehabilitation and may dictate the degree of recovery.

Evidence from human studies support that modulation of cortical function, and specifically, increasing cortical excitation, have clinical implications. For example, removing the afferents of the “good hand” via tourniquet-induced anesthesia, anesthetic block, and constraint induced therapy lead to improved hand function^15,16^. Harnessing the brain’s innate plasticity mechanisms through non-invasive methods such as transcranial magnetic stimulation (TMS) has recently gained interest for use in functional and behavioral research as well as rehabilitation research after brain injury^17-22^ and neurodegenerative diseases^23-28^. Various studies showed promising results using TMS in humans^17,25,29^, primates^30,31^, and rodents^32-36^. Importantly, TMS has shown effectiveness in manipulating transcallosal communication in patients suffering from peripheral injury^37,38^ and alleviating pain associated with injury^39^. TMS has also been shown to increase neuronal excitability and markers associated with plasticity such as brain-derived neurotrophic factor (BDNF)^40-42^, c-*fos*^40^ and Ca^2+^/calmodulin-dependent protein kinase II (CaMKII)^35^. However, it is still not clear when is the optimal and most effective time for TMS intervention (acute, subacute or chronic phase) to take place after injury. Furthermore, TMS stimulates brain tissue nonspecifically which has the potential to induce undesired and unpredictable side effects. Thus, developing neuromodulation strategies to restore normal neural excitability levels with cell-specific precision could lay the groundwork for transforming current clinical practice.

Major advances in molecular and synthetic biology have revolutionized the capability to control cell excitability in living organisms. One of these technologies, magnetic manipulation by the electromagnetic preceptive gene (EPG), allows non-invasive and cell-specific neuromodulation using external magnetic fields. EPG is a protein that is sensitive to electromagnetic fields which was recently identified in the fish *Kryptopterus bicirrhis*^43^. Recent work had demonstrated that calcium imaging in mammalian cells and cultured neurons expressing EPG activated remotely by magnetic fields led to increases in intracellular calcium concentrations, indicative of cellular excitability. Moreover, wireless magnetic activation of EPG in rat motor cortex induced motor evoked responses of the contralateral forelimb *in vivo*. Expressing EPG in S1 contralateral to the injury in rats may provide a way to increase excitation by specifically targeting the excitatory cortical neurons and minimizing off-target affects.

Here we capitalized on multimodal approaches including a battery of behavioral tests, functional MRI (fMRI) and immunohistochemistry to test the effectivity of TMS and EPG-based neuromodulation methods in improving short- and long-term sensory, motor, and cognitive outcomes in a rodent model of peripheral nerve injury.

## Results

### TMS enhances sensorimotor functions

We first tested if non-invasive brain stimulation via rTMS focused on the deprived S1 (contralateral to the denervated forelimb) is an effective strategy to facilitate plasticity and rehabilitation. We performed a battery of behavioral tests to characterize sensorimotor and cognitive functions associated with denervation injury. An illustration depicting the animal model and the different modulation strategies is shown in **Fig 1**. A beam walk test with two different width settings was performed every other week to evaluate sensorimotor functions. The results show that denervated rats that received rTMS treatment every other day for 30 days, starting the day after injury (Den-rTMS-Acute, n=6), showed significantly shorter traverse times and enhanced mobility compared to rats that received rTMS treatment starting 3-weeks after injury (Den-rTMS-Delayed, n=6) and injured rats that received no treatment (Den-No rTMS, n=6). This was true for both the 6.3 cm width (Den-rTMS-Acute, 5.64 ±0.4 s; Den-rTMS-Delayed, 6.35 ±0.4 s; Den-No rTMS, 20.56 ±2.0 s; Control, 5.95 ±0.2 s; p<0.05) and the more challenging, 3.9 cm width beam (Den-rTMS-Acute, 6.29 ±0.4 s; Den-rTMS-Delayed, 7.02 ±0.3 s; Den-No rTMS, 26.4 ±3.4 s; Control, 5.95 ±0.2 s; p<0.05) (**Fig 2.a**). The results show a shortening in the traverse time reflecting an improvement in sensorimotor functions and mobility throughout the course of rTMS treatment. The results demonstrate that at the end of the 4-week rTMS treatment, both the Acute and the Delayed group’s traverse times were shortened and similar to the Control group times; The Den-rTMS-Acute group demonstrated an improvement of 61.3% on the 6.3 cm width challenge beam walk, and a 64.8% on the 3.9 cm challenge beam walk after 4 weeks of treatment. The Den-rTMS-Delayed demonstrated an improvement of 54.7% on the 6.3 cm challenge, and a 55.1% on the 3.9 cm challenge, and the Den-No rTMS showed only a 19% and 37.8% improvement on the 6.3 cm and 3.9 cm challenge, respectively, over the same time frame.

**Fig 1.**
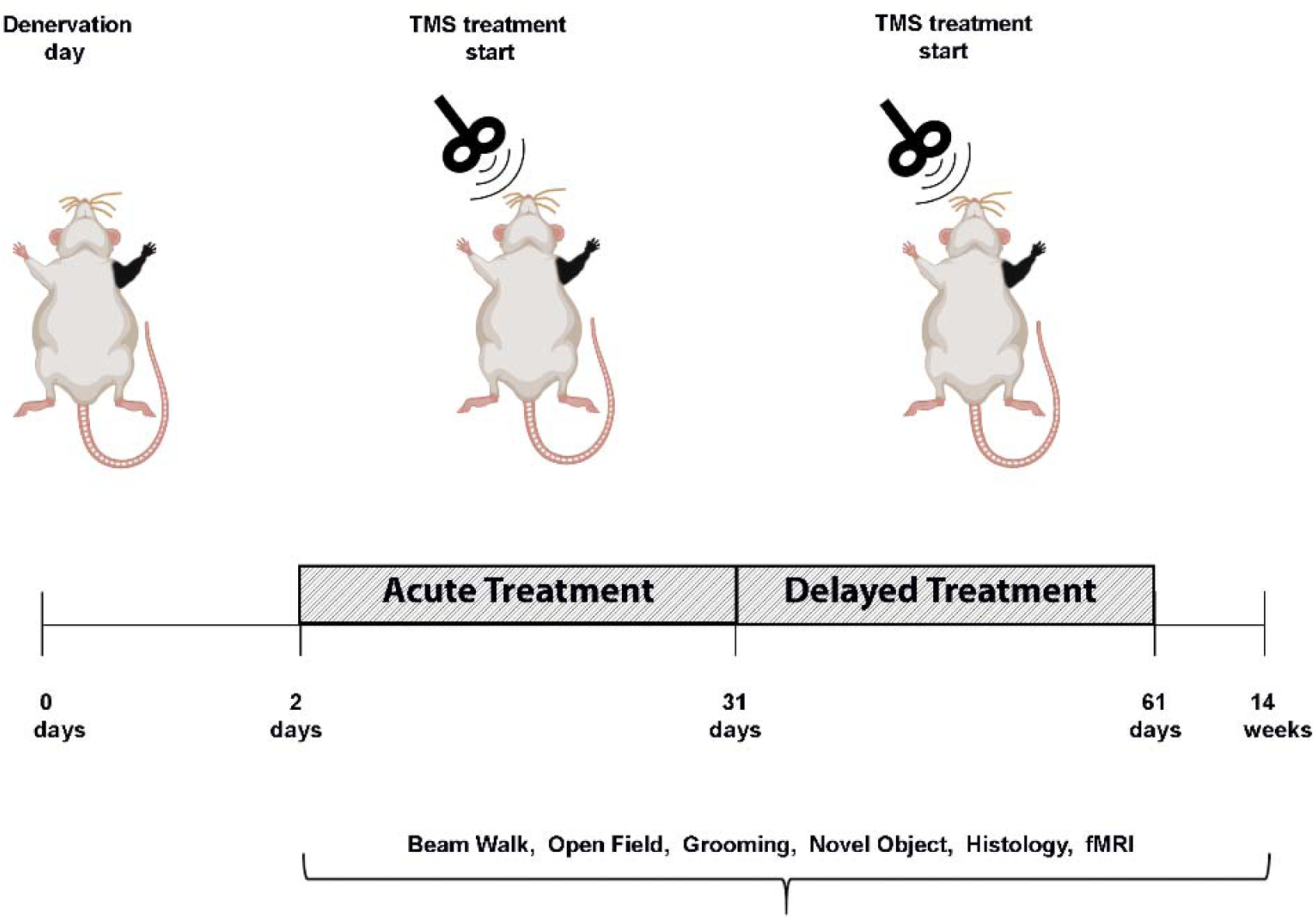
Diagram demonstrating the experimental design of neuromodulation via rTMS. After denervation, rats received rTMS treatment every other day, for 30 days. The intervention began at the acute phase, a day after denervation (Den-rTMS-Acute) or at the sub-acute phase, two weeks following denervation (Den-rTMS-Delayed). The rTMS coil was placed over the left S1, contralateral to the denervated forepaw and delivered 10 min of 10 Hz stimulation. A control group was denervated but did not receive any treatment and an additional control group were not injured and did not receive rTMS treatment.

**Fig 2.**
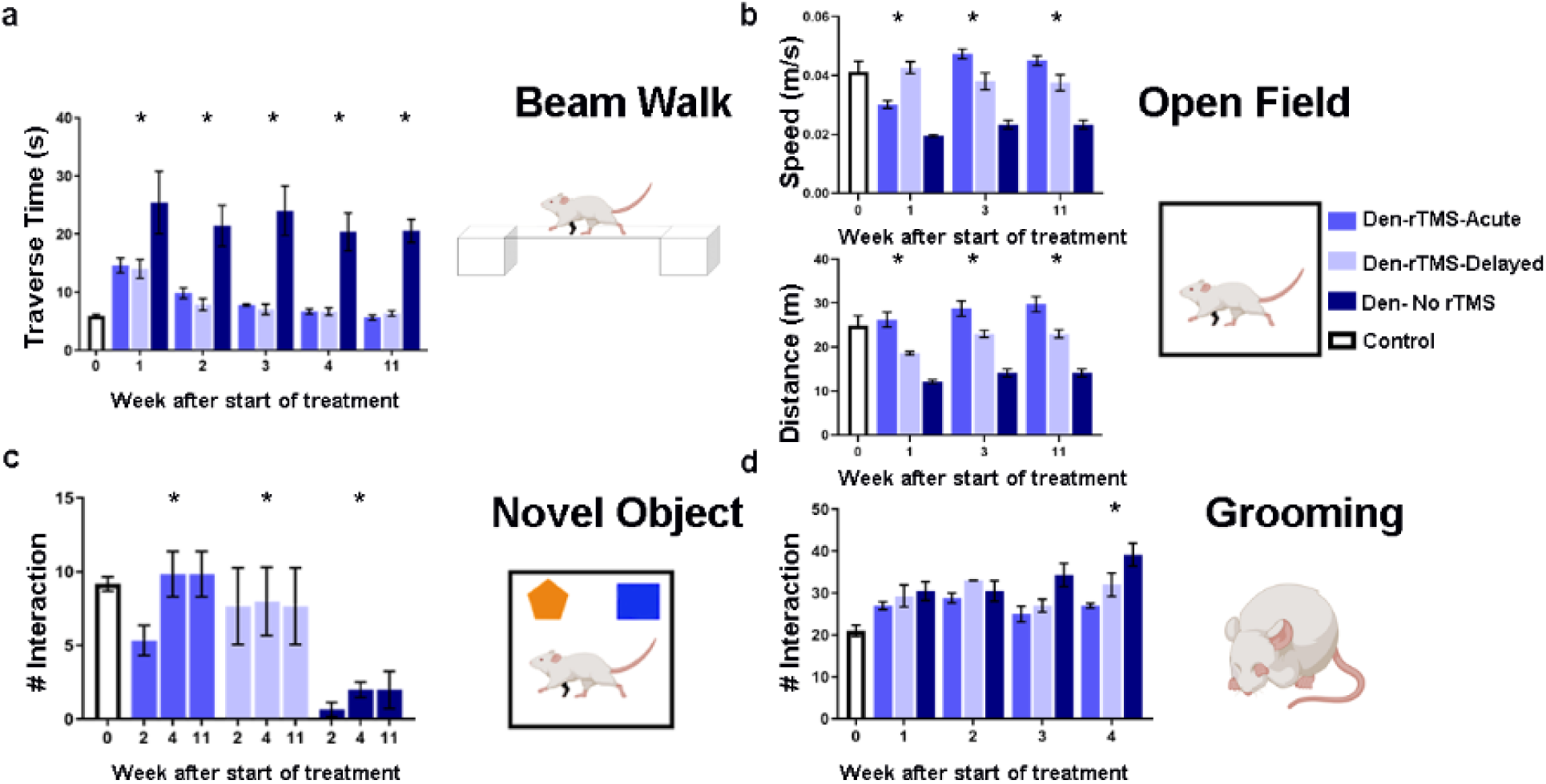
A battery of behavioral tests to assess sensorimotor and cognitive functions was performed before, throughout, and after the rTMS intervention. **a.** Sensorimotor functions were evaluated by the traverse time on a 6.3 cm beam walk. **b.** Sensorimotor and cognitive functions were evaluated by the time and the velocity of movement in the open field arena. **c.** Emotional and cognitive function were evaluated by the time the rats spent exploring new objects in their arena. **d.** Sensory dysfunction and pain associated with denervation was monitored by the number of strokes rats made in the self-grooming test. Results show that rTMS intervention leads to improved long-term mobility and decreased anxiety. Furthermore, rats that received rTMS treatment immediately after denervation (Den-rTMS-Acute) exhibited the greatest improvement compared to Den-rTMS-Delayed and Den-No rTMS (*, p<0.05).

An open field test where rats were placed in a 109 cm × 35.56 cm arena and their movement videotaped with a ceiling camera was performed every other week to assess locomotion and anxiety. During the 10 min test, the denervated rats that received rTMS treatment starting the day after injury showed significant increased in the averaged speed (values at week-4: Den-rTMS-Acute, 0.047±0.0016 m/s; Den-rTMS-Delayed, 0.038±0.0028 m/s; Den-No rTMS, 0.023±0.0014 m/s; Control, 0.041 ±0.003 m/s; p<0.05), and traveled a greater distance compared to the other groups (Den-rTMS-Acute, 28.75±1.75 m; Den-rTMS-Delayed, 23.01±0.8 m; Den-No rTMS, 14.11±0.9 m; Control, 24.88±2.31 m; p<0.05). The results demonstrate that these improvements lasted throughout the rTMS treatment sessions and weeks following the completion of treatment. After 4 weeks of treatment the Den-rTMS-Acute group increased their speed by 56.9%, while the Den-rTMS-Delayed and the Den-No rTMS increased by only 10.9% and 19.6%, respectively (**Fig 2.b**).

A novel object recognition test was carried out to evaluate the rats’ emotional and cognitive functions reflected by the rats’ interest in new objects placed in the open field arena, as was indicated by the number of times the rats approached the object. This test is known to evaluate anxiety and depression levels which are often increased in patients suffering from chronic pain. The results indicated that denervated rats that received rTMS treatment starting the day after injury spent significantly greater time exploring both the familiar (Den-rTMS-Acute, 7.16±0.7 approaches; Den-rTMS-Delayed, 3±1.5 approaches; Den-No rTMS, 3.8±1.7 approaches; Control, 7.6±1.2 approaches; p<0.05) and the novel objects (Den-rTMS-Acute, 9.8±1.5 approaches; Den-rTMS-Delayed, 7.6±2.6 approaches; Den-No rTMS, 2 ± 0.5 approaches; Control, 9.1±0.4 approaches; p<0.05). Rats that received the rTMS treatment immediately after injury have shown the greatest gradual increase in the time they spent exploring the novel object over the course of the treatment regime (Den-rTMS-Acute, 84%, Den-rTMS-Delayed, 66.6%, Den-No rTMS, 33.3%) (**Fig 2.c**).

An additional method to assess sensory dysfunctions and pain associated with injury is monitoring the self-grooming behavior. Over-grooming such as compulsive licking, scratching, and biting on the limbs are often observed in animals suffering from nociceptive pain^44-47^. Rats were placed in a clean cage and videotaped for 20 min, and the number of strokes the rats made during that time was counted. The results show that self-grooming had gradually increased over the weeks after denervation in all denervated rats. However, denervated rats that received rTMS treatment showed significantly less self-grooming compared to denervated rats that did not receive rTMS treatment (Den-rTMS-Acute, 32±2.7 strokes; Den-rTMS-Delayed, 2±0.5 strokes; Den-No rTMS, 39.1±2.7 strokes; Control, 21±1 strokes; p<0.05). Over the time of the study, denervated rats that did not receive treatment had a 28.4% increase in the number of strokes, demonstrating overall increase in pain and discomfort (**Fig 2.d**).

We then sought to determine if the rTMS treatment induced improvements in sensorimotor and cognitive functions that were observed in the behavioral tests also had physiological correlates. We measured whether rTMS treatment led to long-term plasticity and sensorimotor function. Measurements of Blood-Oxygenation-Level-Dependent (BOLD) fMRI responses evoked by tactile stimulation of the non-injured forelimb were performed 12 weeks after denervation, and 8 weeks after the end of rTMS treatment. **Fig 3** shows BOLD fMRI activation *Z* maps of individual Denervated-rTMS-Acute and Denervated-No rTMS rats overlaid on high-resolution anatomical MRI images across S1, as well as the statistics for the groups. The number of activated voxels (General Linear Model statistics with a Z score>2.3, corresponding to p<0.05) in S1 induced by tactile stimulation was calculated for Den-rTMS-Acute (n=5) and Den-No rTMS (n=5) groups. The results demonstrate that rTMS treatment led to significant increase in the number of activated voxels across S1 (Den-rTMS-Acute, 150.8±23.3 voxels; Den-No rTMS, 91.6±7.9; p<0.05) suggesting an increase in neuroplasticity in S1 contralateral to the injured forelimb.

**Fig 3.**
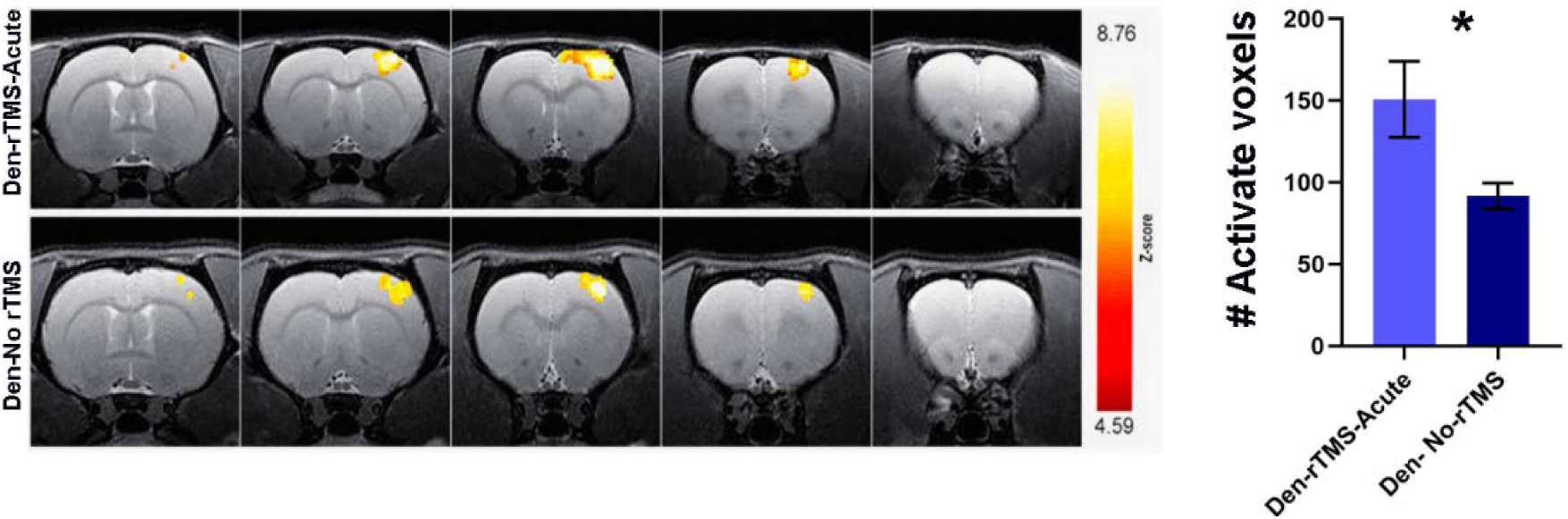
fMRI BOLD responses to intact forepaw stimulation eight weeks after rTMS intervention. Representative BOLD z-score activation maps corresponding to p<0.05, overlaid on high resolution coronal images. **b**. The average number of activated voxels in S1. The significantly greater fMRI activation exhibited by the Den-rTMS-Acute compared to Den-No rTMS suggests enhanced neuroplasticity.

Further immunostaining to identify biomarkers associated with neuroplasticity were performed on 25-µm thick brain slices obtained from rats that were sacrificed 16 weeks after the denervation procedure. We calculated the number of cells and the expression levels of CaMKII, a gene known to be involved in long-term potentiation (LTP). The results showed that rats that received rTMS treatment starting the day after injury exhibited a significantly greater fluorescence intensity of CaMKII (Den-rTMS-Acute, 617.5±60 cells), compared to both rats that received delayed rTMS treatment (Den-rTMS-Delayed, 472±13 cells) and denervated rats that did not receive rTMS treatment (Den-No rTMS, 362±23 cells; ANOVA (p<0.05)). **Fig 4** shows the normalized CaMKII intensity across the deprived S1 (contralateral to denervated forelimb). The non-invasive fMRI and the immunostaining results are consistent with the behavioral tests, and together the results show that rTMS treatment that started at the acute phase after injury led to neuroplasticity and rehabilitation. Furthermore, the results suggest that the rTMS treatment induced long-term neuroplasticity changes that were evident in the behavioral, system, and cellular levels, lasting for months after the treatment has ceased.

**Fig 4.**
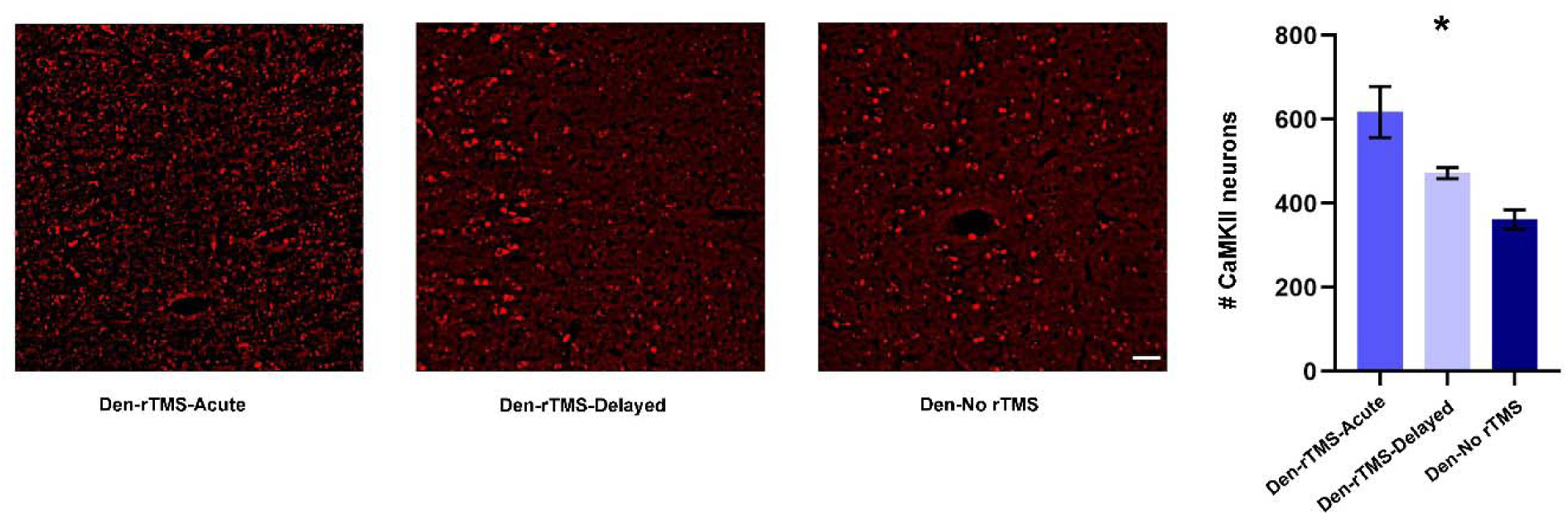
Immunostaining for CaMKII, a marker for neuroplasticity, in S1 contralateral to the denervated forepaw that was subjected to the rTMS intervention. **a.** Microscopy images demonstrate increased fluorescent in S1 neurons in Den-rTMS-Acute and Den-rTMS-Delayed compared to Den-No rTMS. **b.** Quantification of the number of neurons expressing CaMKII (Scale bar = 50 µm).

### Cell-specific neuromodulation via EPG

EPG is a protein that is sensitive to magnetic fields that, upon magnetic activation, increases neural excitability. We tested if expression of EPG in excitatory cortical neurons would restore normal excitation-inhibition balance in deprived S1, which could lead to increased plasticity and rehabilitation.

Right forepaw denervation was performed in 11 rats. One week after the denervation procedure, rats were stereotaxiclly injected with a virus encoding for EPG under the CaMKII promotor (AAV-CaMKII::EPG-GFP). Virus was injected into four different locations covering S1 contralateral to the denervated limb (Den-EPG, n=6). Control rats went through a similar procedure but were injected with virus containing only a GFP marker (Den-Control, n=5). Three weeks after virus injection, and four weeks after denervation, we placed an electromagnet generating a field of 41 mT inside the rat’s skull, directly over S1 expressing EPG, which was contralateral to the denervated limb. A diagram of the experimental paradigm is shown in **Fig 5**. The electromagnet consisted of a ferromagnetic core wound with 2,000 turns of magnet wire, and a simulation demonstrating the magnitude of the magnetic fields experienced by the rat based on the dielectric properties of brain tissue and bone^48^ are shown in **Fig 6.a.** The magnetic stimulation was performed for 16 min once a day, for 30 days, while rats were anesthetized with 2% isoflurane. Immunohistochemistry was performed on brain slices obtained from all of the rats in this study to confirm EPG expression in S1 (**Fig 5**).

**Fig 5.**
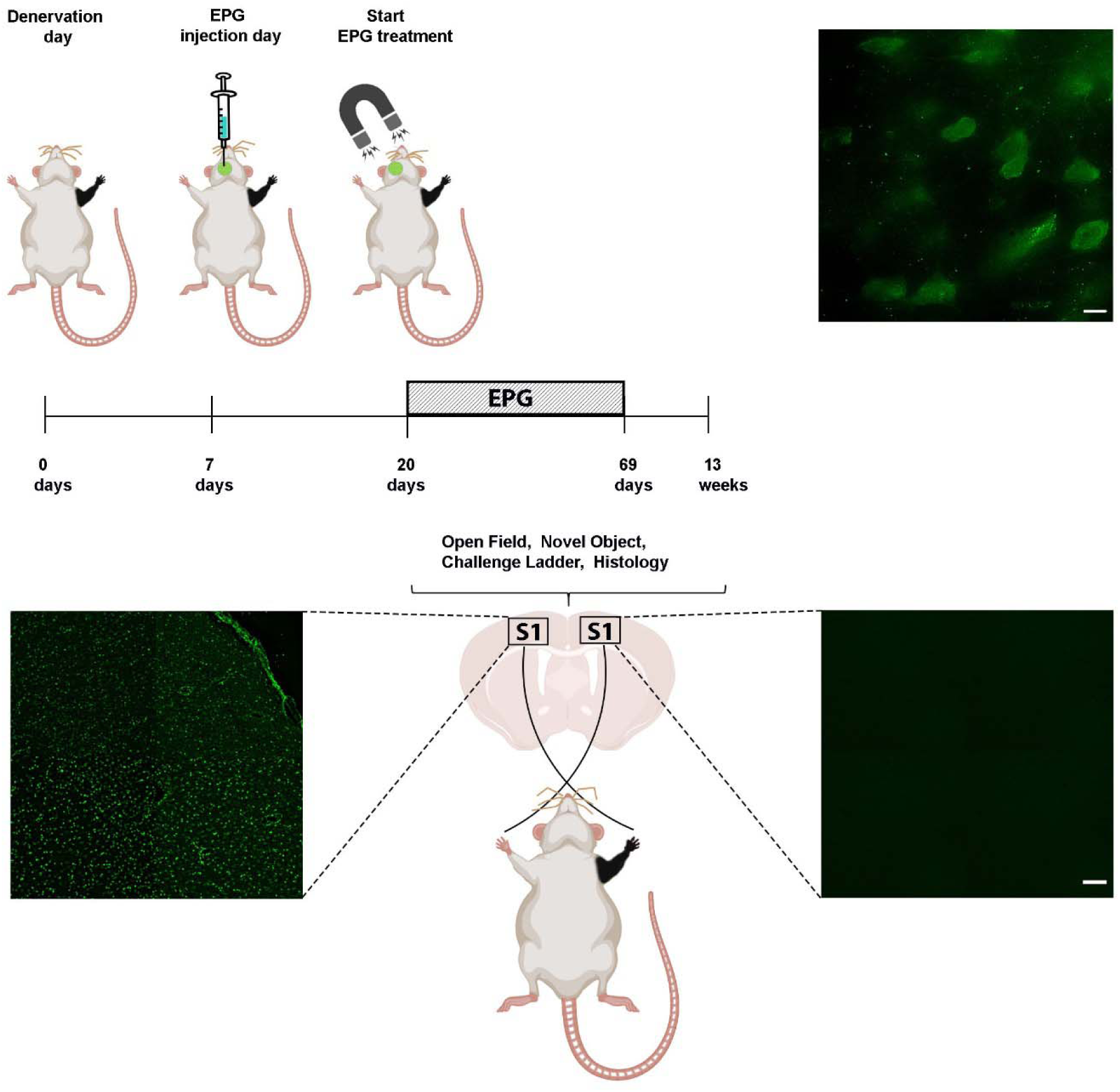
Diagram demonstrating the experimental design of neuromodulation via EPG. Virus encoding to the EPG was stereotaxicly injected into the left S1, contralateral to the denervated forepaw (Den-EPG). Denervated control rats were injected with a virus encoding for a fluorescence protein (Den-Control). An electromagnet was placed over the left S1 starting three weeks following stereotaxic injection. The electromagnet delivered magnetic field stimulation for 16 minutes once a day, for 30 days. Immunostaining images in the primary somatosensory cortex showing EPG expression in fixed brain sections using anti-FLAG antibody in left S1, and right, non-injected S1 with 100X (upper panel, scale bar= 10 µm) and 4X magnification (Scale bar = 50 µm).

**Fig 6.**
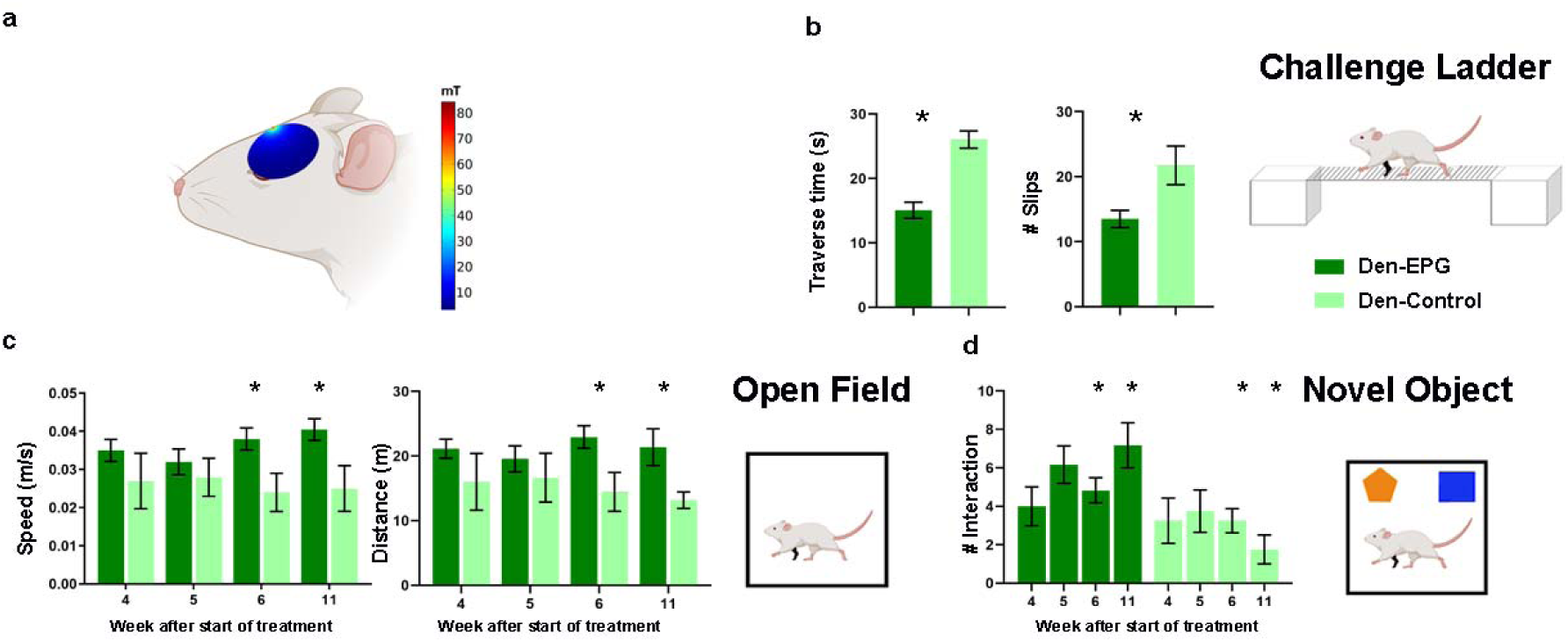
A battery of behavioral tests to assess sensorimotor and cognitive functions was performed throughout and after magnetic activation of EPG. **a**. The magnitude of the magnetic field (mT) in a sagittal plane of a simulated ellipsoidal rat brain. The rat skull and brain were modeled using dielectric properties consistent with human bone and brain tissue. **b.** Sensorimotor functions were evaluated by the traverse time and number of footfalls on a challenge ladder. **c.** Sensorimotor and cognitive functions were evaluated by the time and the velocity of movement in the open field arena. **d.** Emotional and cognitive function were evaluated by the time the rats spent exploring new objects in their arena. The results demonstrate that the Den-EPG exhibited significant and long-term improvement in sensorimotor functions compared to the Den-Control group (*, p<0.05).

A battery of behavioral tests to characterize sensorimotor and cognitive function associated with denervation injury and EPG treatment, was performed throughout the course of the stimulation. Long-term improvement in sensorimotor functions and mobility was evaluated using the challenge ladder test, whereas the travers time and the number of slips were determined by laser sensors. This test was performed 12 weeks following denervation, 8 weeks after EPG injection, and 4 weeks after the EPG magnetic stimulation treatment ended. The results show that Den-EPG rats had crossed the ladder in a significantly shorter time (Den-EPG, 15.08±1.2 s; Den-Control, 26.05±1.3 s; p<0.05) and exhibited fewer slips (Den-EPG, 13.5±1.3 s; Den-Control, 21.75±2.9 s; p<0.05) (**Fig 6.b**).

Open field was performed once a week throughout the EPG magnetic treatment. The results demonstrated that within three weeks after starting the magnetic stimulation, the denervated-EPG rats showed significant increases in speed (values at week-4: Den-EPG, 0.04±0.002 m/s; Den-Control, 0.02±0.006 m/s; p<0.05), and traveled a greater distance compared to the control group (Den-EPG, 22.94±1.7 m; Den-Control, 14.45±3 m; p<0.05) (**Fig 6.c**). These improvements lasted for a month after the magnetic stimulation treatments ended, suggesting that the EPG manipulation induced long-term neuroplasticity changes in S1 circuitry.

Den-EPG rats also demonstrated increased interest in new objects during the novel object recognition test. Significant increase of the time they spent exploring the new object, compared to controls, was observed four weeks after the magnetic stimulation treatment ended (Den-EPG, 7.16±1.1 approaches, Den-Control, 1.75±0.7 approaches; p<0.05) (**Fig 6.d**). Overall, the results show that EPG neuromodulation in denervated rats led to substantial improvement in sensorimotor function and rehabilitation.

## Discussion

Our study demonstrates that neuromodulation in the days and the weeks following peripheral nerve injury leads to short-term and long-term plasticity and neurorehabilitation. Daily neuromodulation regimes with rTMS have shown to improve sensory, motor, and an overall well-being of the injured rats in a battery of behavioral and imaging tests that were performed up to 8 weeks after the rTMS treatment ended. The results suggest that an immediate, acute, rTMS intervention may be more effective compared to rTMS treatments that begins in the sub-acute phase. This builds on a growing bulk of evidence demonstrating that peripheral nerve injury leads to immediate changes in neural function that may dictate the degree of future rehabilitation^12,14,37,49-51^. Immediate changes in both spontaneous and evoked neural activity have been also demonstrated in models of spinal cord injury^52^. Indeed, and early intervention of rTMS therapy in a rodent model of spinal cord injury has also shown to be more effective compared to later-stage intervention^34^. Nevertheless, delayed rTMS stimulation also led to behavioral improvement compared to rats that did not receive any treatment. Thus, post-injury rTMS treatments may be tailored to benefit patients in the acute, sub-acute, and even chronic phases.

Neuromodulation by rTMS may provide an effective, accessible, relatively inexpensive and completely non-invasive approach to attenuate pain associated with peripheral nerve injury and improve sensorimotor outcomes. However, the shortcoming of this technique includes non-specific stimulation that could lead to undesired side effects, and on rare occasions, it can evoke seizures. Thus, there are ongoing efforts to develop minimally invasive therapeutic strategies that will diminish non-specific activation but will allow temporal precision. Tools such as optogenetics and chemogenetics have the advantages of cell type specificity and superior spatial and temporal resolution compared to prior neuromodulation methods. Indeed, we have previously shown that neuromodulation via optogenetics approaches was successful in restoring cortical excitation-inhibition balance in the weeks following the peripheral nerve injury^13^. Specifically, light activation of halorhodopsin in the healthy cortex combined with forepaw stimulation lead to increase of excitatory neuronal activity in the deprived somatosensory cortex of peripheral nerve injured rats. However, one of the drawbacks of this technology is the requirement to deliver the light directly into the target neural population. Here we tested if neuromodulation via the magnetic sensitive protein EPG, which provides cell and temporal specificity while being activated remotely via non-invasive electromagnetic fields^53^, can be utilized to restore cortical excitability and achieve similar sensorimotor outcomes compared to rTMS. The results demonstrate that daily magnetic activation of EPG improved sensory, motor, and an overall well-being of the injured rats in a battery of behavioral tests that were performed up to 4 weeks after the EPG treatment ended.

Growing amounts of evidence from human and animal studies are establishing neuromodulation as an effective mechanism to strengthen and promote cortical functions^54^. The behavioral results indicate that both rTMS and EPG treatment have led to considerable improvement in sensorimotor functions. Overall, it appears that rTMS treatment delivered at the acute phase had the most significant impact on physiological functions compared to rTMS delivered after a few weeks after injury, and EPG neuromodulation that also began a few weeks after the injury. Thus, it is manifested that an immediate intervention may be the most effective in reshaping and guiding post-injury plasticity. In addition, neuromodulation via EPG is a new and upcoming technology and efforts are being made towards discovering the molecular structure and the signal transduction basis of this phenomenon. It is also anticipated that utilizing synthetic and molecular biology approaches as well as improving in the hardware will make the EPG function more robust. The EPG technology complements other neuromodulation methods and expands the current toolbox for basic and translational research.

## Acknowledgments

This work was supported by National Institutes of Health grants R01NS072171, R01NS098231 and R01NS104306.

## Author contribution

CC and GP designed all experiments and wrote the manuscript. CC, MZ, AM, CQ performed experiments. CC, GP, and MZ analysed the data. RA and LU performed the device fabrication, magnetic measurements, and related analysis. CC, RA, MZ, AM, CQ, LU, AAG, and GP were involved in the discussions and preparation of the manuscript and approved the final manuscript.

## Methods

### Animals

All animal procedures were conducted in accordance with the NIH *Guide for the Care and Use of Laboratory Animals* and approved by the Michigan State University Animal Care and Use Committee. Thirty-five Sprague-Dawley rats (19 male and 16 female) were provided with food and water *ad libitum* and housed in a room with a reverse cycle.

### Surgeries and Stimulation

Forepaw denervation was performed on 29 rats weighing 80-90 g. Rats were anesthetized with 2% isoflurane which was delivered through a nose cone. Skin incision was made on the right forepaw, and the radial, median and ulnar nerves were cut, and a 5 mm gap was made in each one. The incision was closed with silk sutures and tissue glue. Tramadol (0.1 mg/300 mg) was administrated orally for 5 days after the injury.

The TMS system was equipped with a figure eight, 25 mm custom rodent coil (Magstim, Rapid2) that was placed over the left hemisphere directly on the head. TMS was delivered once a day, for 31 days with the following settings: 4 s cycles of 10 Hz stimuli, 26 s interval, and 7 cycles (total of 280 pulses per day, 1680 total stimuli). During the stimulation, rats were anesthetized with 2% isoflurane. Denervated control rats that did not receive the TMS treatments were subjected to the same daily anesthesia protocol for the same length of time.

Eleven rats received stereotaxic injection a week following the denervation surgery: 6 of them were injected with virus encoding for EPG under CaMKII promoter (pAAV2-CaMKII::EPG-IRES-hrGFP), and 5 with virus encoding only to GFP (pAAV2-CaMKII::IRES-hrGFP). Rats were anesthetized with 2% isoflurane which was delivered through a nose cone and secured in a stereotaxic frame. The microinjection needle was placed in four locations in the left primary somatosensory cortex (S1) area: AP: +0.2 mm and +0.3, ML:−3.8 mm and −3.2. A volume of 1 μL of virus was injected in each location starting at a depth of 1.2 mm and retracting the needle up to a depth of 0.8 mm.

Rats were divided into the following six groups: **1.** Denervated rats that started receiving rTMS 48 hours following denervation (Den-rTMS-Acute, n=6). **2.** Denervated rats that started receiving TMS 3 weeks following denervation (Den-rTMS-Delayed, n=6). **3.** Denervated rats not receiving TMS (Den-No rTMS, n=6). **4.** Non-denervated not receiving TMS (Control, n=6). **5.** Denervated rats injected with virus containing EPG in S1 contralateral to denervated limb (Den-EPG, n=6). **6.** Denervated rats injected with virus containing only GFP in S1 contralateral to denervated limb (Den-Control, n=5).

### Behavioral Assessments

A comprehensive battery of behavioral tests to assess sensory, motor, and cognitive functions was performed over 30 days since the beginning of TMS therapy. Grooming: Rats were placed separately in a clean cage (43.62cm (L) × 22.86cm (W) × 20.32cm (H)) with food and water. Grooming was recorded for 20 min. The first minute was considered habituation period, and the rest of the 19 min were analyzed. The number of interactions on each part of the chain grooming actions was counted for each individual. This test was performed once a week.

#### Open Field

The open field was carried out in an arena with the following dimensions (L) 109 cm × (W) 35.56 cm × 142.24 cm (H) (San Diego Instruments). During the session, the open field was isolated from the observer, and the light intensity was maintained stable. Movements were recorded by a ceiling mounted camera for 10 min. The freezing time, total distance and averaged velocity were analyzed by an automated tracking system (ANY-Maze software, San Diego, USA). After each session the arena was cleaned with 70% ethanol. This test was performed every two weeks (for TMS treated rats), and once a week (EPG rats).

#### Novel Object Recognition

Rats were placed in the open field arena. In the first stage, rats were acclimating to the environment (5 min). In the second stage two identical objects were placed in the arena and the rat got familiarized with them (5 min). In the third stage we replaced one of the objects for a new and unfamiliar object (5 min). The time spent exploring the novel object was analyzed by automated tracking (ANY-maze software). This test was performed every two weeks (for TMS treated rats), and once a week (EPG rats).

#### Beam Walk Test

TMS-treated rats were placed on one end of 114.3 cm-long suspended, narrow wooden beam. Two different widths were tested: 6.3 cm and 3 cm. The traverse time from one end to the other was measured. Three training sessions were performed for each animal once a week. For the denervated-TMS group, the rats started to walk on the 6.3 cm width beam and then were challenged on the 3 cm wide beam.

#### Challenge Ladder

Den-EPG rats crossed a 114 cm-long horizontal suspended ladder with rungs spaced 1.3 cm apart (San Diego Instruments, USA). The traverse time and the number of failures to place the paw correctly on the ladder were observed. Two training sessions were performed for each animal on test days, with this test being performed only once.

### Functional MRI

fMRI activity was assessed in denervated rats that received TMS (TMS acute, n=5) and rats that did not receive the TMS (denervated no-TMS, n=5). Rats were anesthetized with dexmedetomidine (0.1 mg/kg/h, SC) which is known to preserve neurovascular coupling. Rats were then placed in a 7 T/16 cm horizontal bore small-animal scanner (Bruker BioSpin, Rheinstetten, Germany). A 72-mm quadrature volume coil and a 15-mm-diameter surface coil were used to transmit and receive magnetic resonance signals, respectively. Respiration rate, heart rate, and partial pressure of oxygen were continuously monitored throughout fMRI measurements (Starr Life Sciences, Pennsylvania, USA). For fMRI, FID-EPI was used with a resolution of 150 × 150 × 1000 μm. Five, 1 mm thick coronal slices covering the primary somatosensory cortex (S1) were acquired (effective echo time (TE), 16 ms; repetition time (TR), 1000 ms; bandwidth, 333 KHz; field of view (FOV), 3.5 × 3.5 cm; matrix size, 128 × 128). A T2-weighted TurboRARE sequence was used to acquire high-resolution anatomical images (TE, 33 ms; TR, 2500 ms; bandwidth, 250 KHz; FOV, 3.5 × 3.5 cm; matrix size, 256 × 256) corresponding to the fMRI measurements. Two needle electrodes were inserted into the left and right forepaws to deliver electrical stimulation. Electrical stimulation was applied in two 40 s trains (3 Hz, 0.4 mA, and 0.4 ms). fMRI analysis was performed using SPM fMRat software (SPM, University College London, UK). Activation maps were obtained using the general linear model. The experimental design was rest <stimulate, T-contrast Thresholded:FWE (Family Wise Error) and P<0.05, Extent threshold k=2, Z-score statistics were used with a threshold of Z>2.3.

### Electromagnetic stimulation

The electromagnet used to deliver the magnetic stimulation to the rats’ brains consisted of a ferromagnetic Iron-Nickel core wound with 2,000 turns of 30 AWG magnet wire. Iron core had dimensions of 16.29 cm in length and a diameter of 1.05 cm. A 45 Degree angle was cut into each end, so that both tips of the core came to a point. During stimulation, 5 V was applied to across the connections of the magnet and a current of about 390 mA flowed through the coil to generate the magnetic field. An external digital signal was used to turn the magnet on and off. Measurements of the magnet at varying distances from the core demonstrated that a magnetic field value of 41 mT was generated just in front of the core.

Finite element analysis was performed using ComSol Multiphysics to simulate the magnetic field stimulation delivered to the rat brain. An electromagnet with 2,000 turns wound around a ferromagnetic Iron-Nickel core with a length of 16.29 cm, a diameter of 1.05 cm, and a relative permeability of 100,000 was used to stimulate the rat brain. A current of 390 mA was passed through the simulated coils and the tip of the core was placed 0.5 mm from the surface of the skull. The rat skull and brain were modeled by concentric ellipses, the larger of which having dimensions 21 mm × 11 mm × 16 mm, with a skull thickness of 0.7 mm used^48^. Dielectric properties for human bone and brain tissue were used to represent bone and brain tissue in the rat, specifically a relative permittivity of 1.53 × 10^3^ and 6.10 × 10^4^, a relative permeability of 1 and 1, and a conductivity of 2.03 × 10^−2^ S/m and 1.06 × 10^−1^ S/m respectively^48^.

### Immunochemistry of Brain slices

Rats were perfused with 0.1 M phosphate buffer saline solution (PBS) in pH 7.4 followed by ice cold 4% paraformaldehyde solution and the brains were removed. Brains were sliced on a cryostat to obtain 20 µm thick sections. Sections were incubated overnight with primary antibodies to detect CaMKII (anti-CaMKII rabbit, Abcam #ab52476); FLAG (anti-flag mouse antibody, Abcam #ab49763), and GFP (anti-GFP chicken polyclonal antibody, Abcam #ab13970). Sections were incubated for 3 h at room temperature with secondary antibodies, processed with ProLonng Gold antifade reagent with DAPI (Thermo Fischer Scientific 2078923) and then imaged on the DeltaVision microscope. ImageJ was used for analysis.

### Statistics

The number of rats in each group was determine by a power of 0.93 assuming an effect of 1 standard deviation while minimizing the number of rats in each group. All the results and figures show mean ± standard error of mean (SEM). Two-tailed Student’s *t*-test and an ANOVA were used when appropriate.

